# From flocks to swarms: rewriting the rules of collective cell migration

**DOI:** 10.64898/2026.09.29.753501

**Authors:** Nicholas R. Martin, Nolan Brown, Roman Gaidarov, Isabella Wischik, Ariel Amir, Ajay Gopinathan, Orion D. Weiner

## Abstract

From flocking birds to swarming insects, collective motion enables groups to accomplish tasks beyond the capabilities of individuals. These behaviors emerge from local interactions among group members, but the interaction rules that organize collectives at the cellular scale remain poorly understood, in part because they are difficult to manipulate experimentally. Here we develop a cell-machine interface that enables us to rewrite the rules of collective migration. Cell-cell interaction rules are specified in software and implemented in living cells through real-time computer vision and closed-loop optogenetic control. Using a single engineered cell line, we program cells to self-organize into flocks, to propagate regenerative signal relays that recapitulate the interaction logic of immune-cell swarms, and to collectively navigate guidance cues that individual cells cannot. Our work shows that altering simple rules of cell-cell interaction is sufficient to generate new emergent collective capabilities. This approach provides an experimental framework for discovering the organizational principles of multicellular coordination and designing synthetic collectives with capabilities beyond those of their individual members.

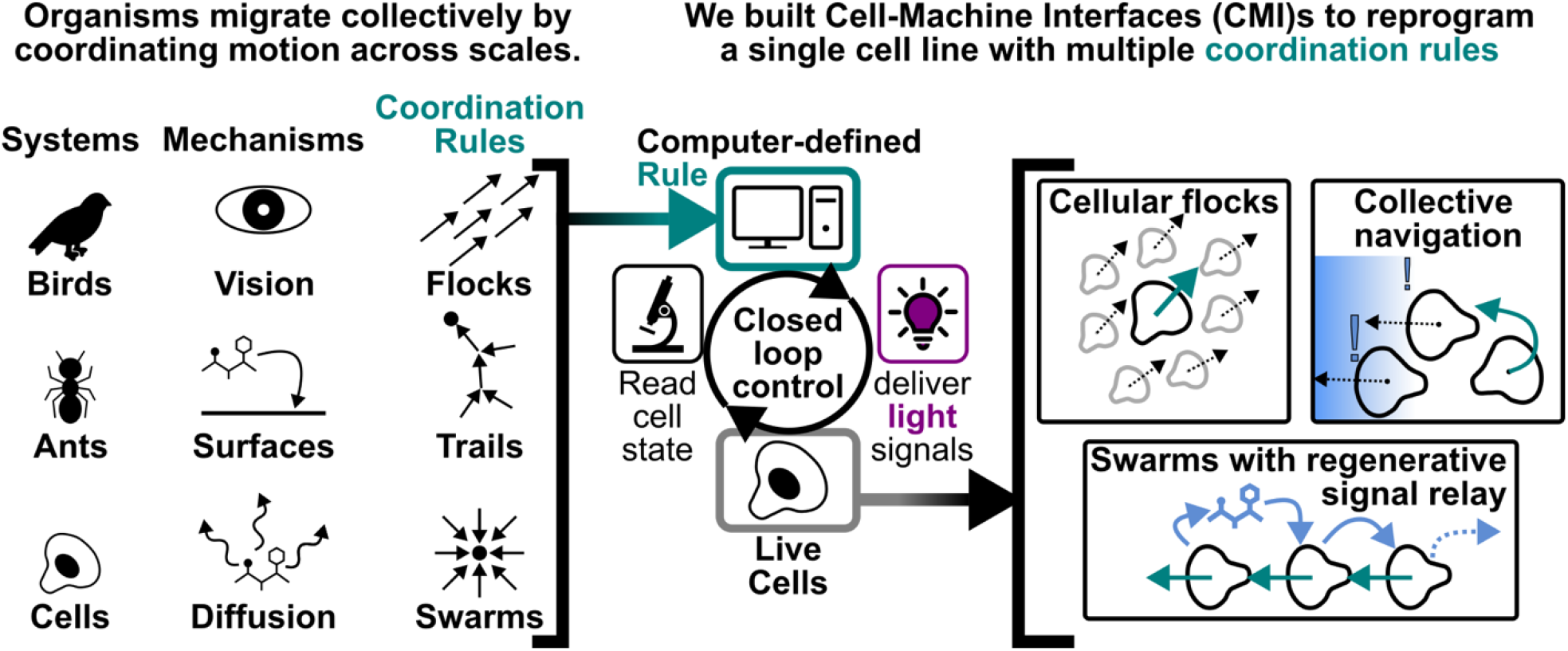

## Introduction

Collective migration enables groups to accomplish tasks that individuals cannot achieve on their own. At the organismal scale, flocking birds align their motion to evade predators^1^ and ants reinforce pheromone trails to organize foraging^2,3^. At the cellular scale, collective migration occurs during immune-cell recruitment^4–6^, wound healing^7^, and developmen^8^. Observational studies^9,10^, perturbation of key information carriers^6,11^, and mathematical modeling^12,13^ have all provided insight into how local interactions among individuals generate population-scale collective migration. However, understanding the general principles across collective systems remains challenging they operate across different spatiotemporal scales, use distinct modes of communication, and perform different collective tasks. Here, we undertook a complementary approach of synthetically reprogramming the local rules of cell-cell interaction that drive collective migration.

Synthetic biology is a powerful approach for testing whether we understand the logic of a given system well enough to build it from scratch. This strategy has been used to understand the basis of biological oscillators^14,15^, cell state transitions^16^, and multicellular pattern formation^17^. We sought to make collective migration similarly programmable by rewriting the local rules of cell-cell interaction. Collective migration offers a particularly rich repertoire of local coordination rules, many of which have been formalized in theoretical models, including neighbor alignment, trail following, and diffusible signaling relays. Yet experimentally testing how distinct local interaction rules give rise to collective motion remains challenging. Here we develop a new approach to systematically define and test these rules. Our strategy enables precise definition of cell-cell interaction rules, including forms of interaction that would be difficult or impossible to implement using standard molecular approaches. Because these rules are encoded in software, their logic can be rapidly exchanged while keeping the cellular chassis constant.

Typical synthetic approaches reprogram cellular processes by introducing new proteins, requiring the cell line to be genetically rebuilt for each new circuit^18^ which can result in a lag of months or more between design-build-test cycles. Furthermore, many cell processes are not sufficiently modular to enable conventional synthetic control with defined input/output logic. Here, we take a different approach in which the input/output rules are encoded by a computer but implemented in cells via closed feedback optogenetic control. This strategy transforms the process of reprogramming cells from molecular design and validation into updating a few lines of code. This enables precise control of input/output logic, greatly expands the range circuits that can be constructed, and reduces the time between iterations from months to minutes. Our work goes beyond classical optogenetics, which typically perturbs cellular processes to probe their existing input/output logic. By combining optogenetic control with real-time computer vision, we impose new input-output relations, effectively rewriting cell decision-making in real time. Our work is related to but goes beyond pure computational simulations because we are exploring parameter space in the rich and complex setting of actual cells that that aren’t understood in sufficient detail to purely model in silico.

Rewriting the rules of cell-cell interaction with a CMI requires three components: a mechanism for reading out cell state, a means for controlling cell behavior, and a closed-loop architecture that defines the input-output relation between them (**Fig. 1A**). Here, we read out cell migration using automated computer-vision and control cell migration using subcellular optogenetic manipulation of the migration machinery. The CMI links these measurements and perturbations via user-defined input-output logic that enables us to specify new rules of cell-cell interaction. CMI rules connect past cell behavior to future optogenetic stimulation by algorithmically defining how each cell’s state should influence the stimulation of other cells (**Fig. 1B**). Cells complete the feedback loop by changing their state in response to light (**Fig1. A-B**). We used optogenetic phosphoinositide 3-kinase signaling (opto-PI3K^19^) to control the migration of individual cells based on the behavior of their neighbors (**Fig. 1C, Supplemental Video 1**). These CMI-based cell-cell interaction rules suffice to make cells flock like birds (**Fig. 2, Supplemental Videos 2-4**), collectively solve guidance tasks that are beyond the ability of individual cells acting alone (**Fig. 3, Supplemental Video 5**), and swarm like neutrophils (**Fig. 4 Supplemental Videos 6-7**). Because these rules are written in computer software instead of cellular hardware, we can implement these diverse interaction rules without needing to genetically modify the cell for each iteration.

**Figure 1:**
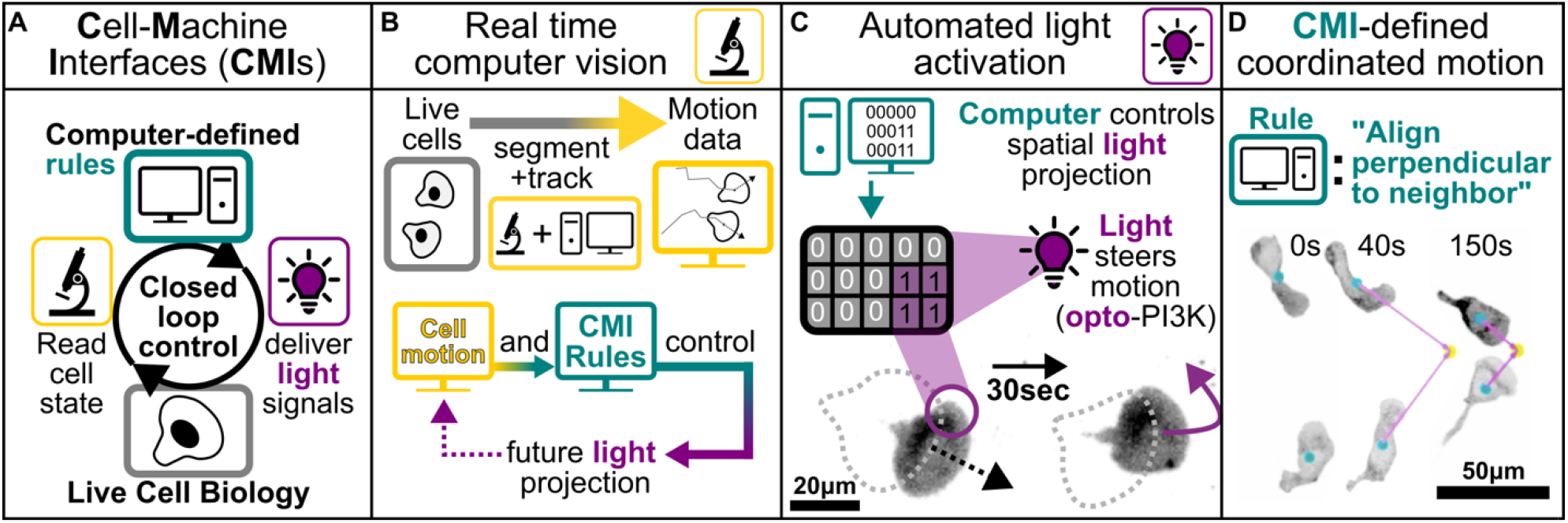
Rewriting the Rules of Cell-Cell Interaction with a Cell-Machine Interface. **A)** The Cell-Machine Interface (CMI) enables computer-defined cell-cell interaction rules by measuring the behavior of one cell and using it to control the behavior of other cells through a closed feedback loop. **B)** The input for the CMI is the state of cells in the field (such as migration trajectory), assayed via real-time computer vision. The output of the CMI is optogenetic control of cell behavior, such as migration orientation. With the CMI, we can define rules that link the inputs with the outputs to enable different rules of cell-cell interaction like neighbor collision (Figure 1D) or alignment (Figure 2). **C)** Subcellular light projection converts output from the CMI rule into changes in live cell motion via an engineered light-sensitive migration regulator (opto-PI3K^19^). **D)** When the CMI aligns the motion of neighboring cells perpendicular to each other, collisions emerge without specifying either the collision point or the paths cells take to reach it.

**Figure 2:**
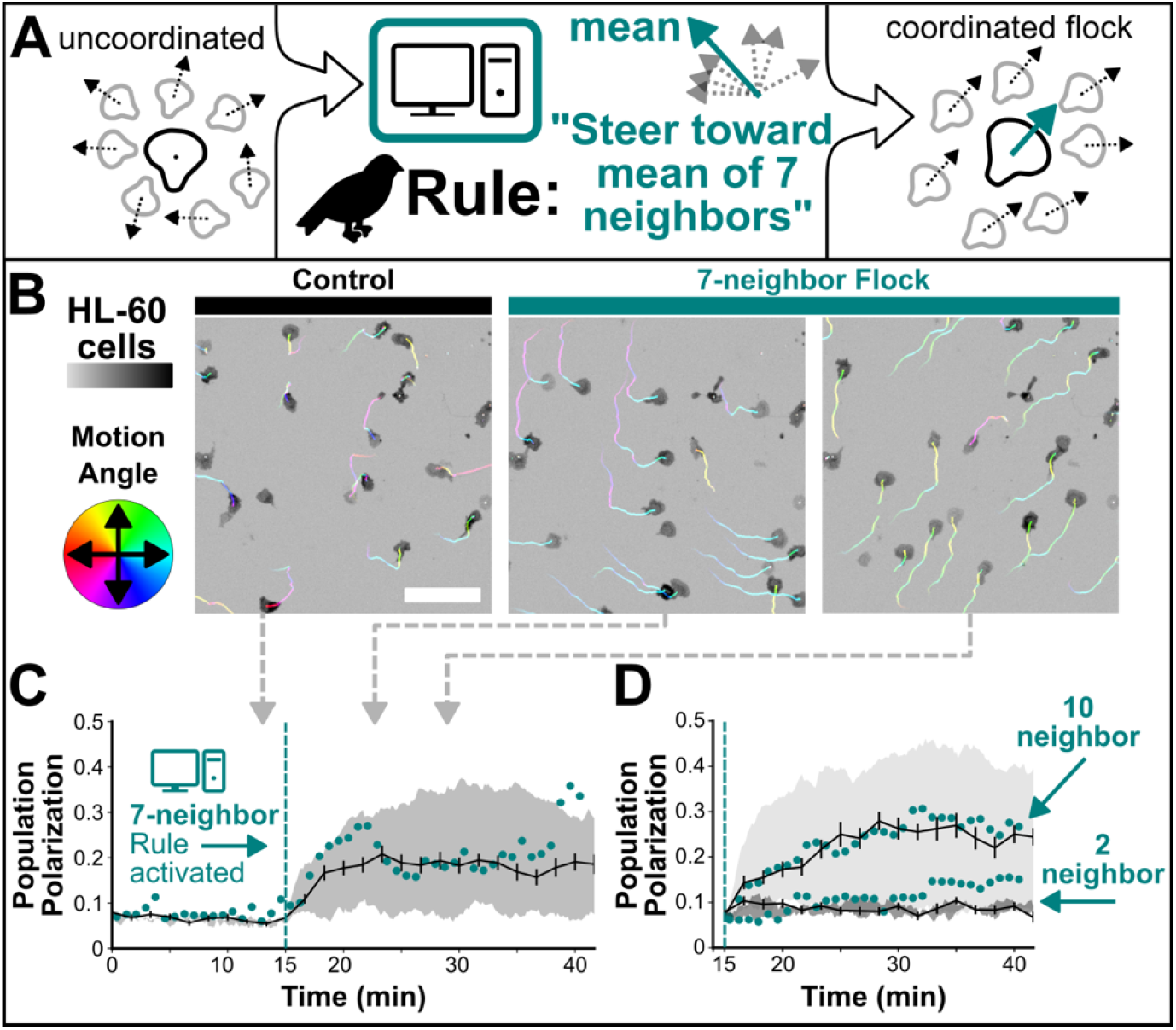
Bird Flocking Rules Program Collective Self-Organization in Migrating Cells. Here we sought to reprogram living cells to interact based on the iconic rules of bird flocking. **A)** Bird flocking is thought to arise from individuals aligning their motion with their seven closest neighbors. **B)** Implementing this rule with the CMI caused normally uncoordinated neutrophil-like HL-60 cells (left) to self-organize into flocks (middle, right). **C)** Mean polarization of simulated (black line) and experimental (cyan points) populations over time before (left of dashed line) and after 7-neighbor flocking CMI rule activation. **D)** In simulations of bird flocking, the number of neighbors over which each individual averages is a key parameter that controls collective behavior. This parameter cannot be readily manipulated in living birds but can be rapidly and precisely manipulated in living cells with a CMI and produces the expected quantitative changes in Polarization. In **B** track segments are color-coded by angle of motion and scale bar is 100μm, all images to scale. Gray arrows point to corresponding time point in **C**. Error bars in **C** are SEM of total polarization from simulations, and shaded region is simulation +/− one standard deviation of the angular velocity parameter (see method details).

**Figure 3:**
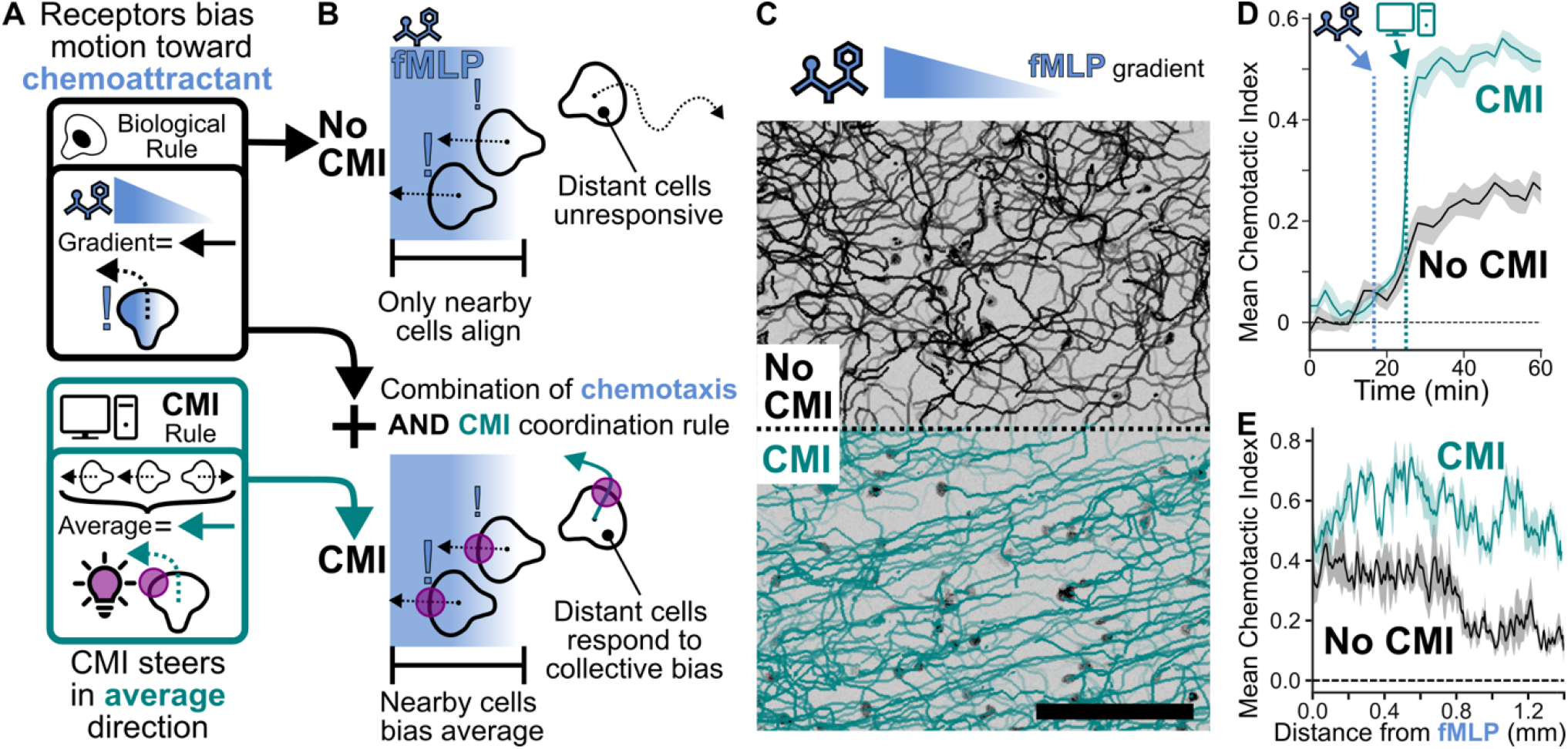
Global coordination enhances collective chemotactic guidance. We next asked whether coordination enables cellular collectives to solve real biological problems. **A)** HL-60 cell sense and respond to chemotactic gradients as individuals without significant multicellular coordination (upper panel). We introduced a global coordination rule to steer cells toward the average motion of the population (lower panel). **B)** Cells that interpret gradients as individuals can only align with chemoattractants when close to the source (upper panel). We hypothesized that synthetically coordinated cells would extend their range of sensitivity (lower panel). **C)** Globally coordinating motion via CMI (bottom) enables enhanced gradient sensing compared to cells lacking this coordination (top); both groups of cells are in same field of view, but CMI is engaged only in bottom cells. **D)** Mean chemotactic index over time from three biological replicates +/− SEM. Vertical lines denote onset of chemotactic gradient and CMI rule, respectively. CMI-enabled global cell coordination significantly improves gradient sensing for the population. **E)** Mean chemotactic index as a function of distance from the left edge of the chemotactic gradient. The CMI led to high gradient alignment both close to and far from the chemoattractant source, whereas cells lacking the CMI had poor gradient alignment far from the source. Binned data from plateau in D (45-60min) +/− SEM across 3 biological replicates. Bar in **C** is 500 μm.

**Figure 4:**
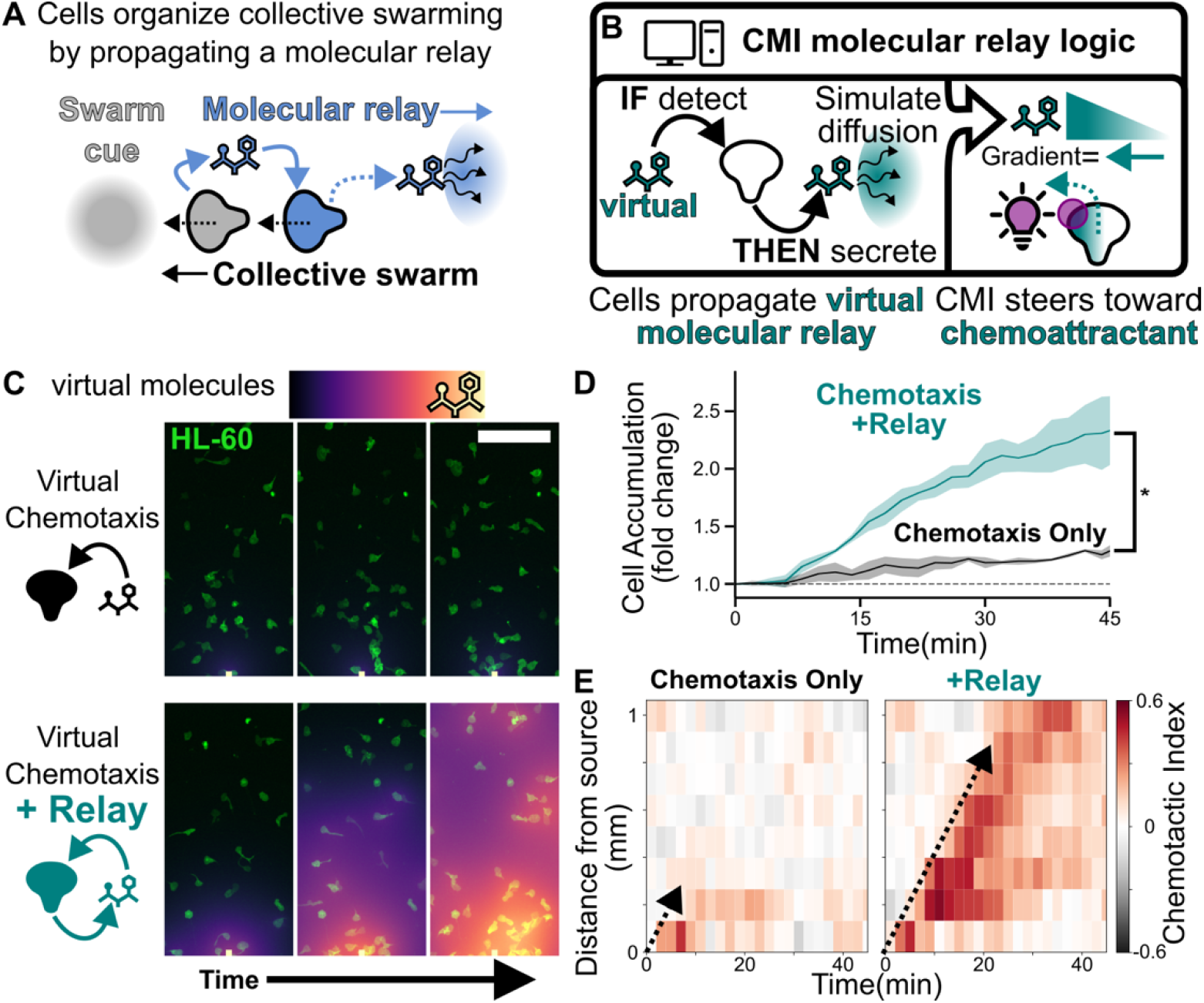
Synthetic signal relays propagate guidance information to enhance long-range recruitment. In our previous experiment (**Fig. 3**), cells globally coordinate. Here we test a more local coordination inspired by the rules of neutrophil swarming. **A)** During neutrophil swarming, pioneer cells that arrive at sites of injury and infection recruit additional cells by releasing a secondary chemoattractant. This signal attracts nearby cells and induces them to secrete chemoattractant themselves, generating a rapidly propagating self-amplifying signal relay. **B)** We used the CMI to implement virtual chemotaxis and relay rules in living cells. The computer simulates the production and diffusion of a virtual chemoattractant, while living cells sense and respond to the resulting gradients through CMI-directed migration. Cells exposed to sufficient virtual chemoattractant become secondary sources, creating a regenerative relay. **C)** Compared to cells using the chemotaxis rule alone (upper), populations also programmed with the relay rule (lower) propagated a wave of the virtual chemoattractant across the field of view. **D)** The relay significantly enhanced cell recruitment to the target. **E)** Because guidance information is transmitted from cell-to-cell, this results in a propagated wave of cell alignment with the target (dashed arrow, right), as seen in native neutrophil swarms. Cells lacking the relay only align close to the target. The Scale bar in **C** is 200μm, images are approximately from 8,16, and 30 minutes. In **D**, shaded region is mean fold change from initial starting density +/− SEM from three technical replicates. Student’s t-test: p=0.03.

We first demonstrated CMI-mediated control of cellular interactions in the simplest possible multicellular context: two cells programmed to collide with one another. A CMI rule that steered each cell perpendicular to the motion of its nearest neighbor produced collisions in which each cell adjusted to the speed, persistence, and starting trajectory of the other cell (**Fig. 1D, Supplemental Video 1**). Importantly, the CMI did not specify the collision location, or the path cells take to reach it. Instead, just as natural collectives coordinate through local interactions, these collisions emerge from the motion of two individual cells interacting through a locally-defined rule—in this case, one implemented through a computer. Because future optogenetic stimuli depend on both the CMI interaction rule and on past cell behavior, the control is intrinsically reciprocal. The computer shapes cell behavior, and the cell behavior controls the computer’s subsequent action.

We next sought to rewrite the rules of cell-cell interaction using rules from the iconic collective behavior of bird flocking. Bird flocking is thought to arise by each bird aligning its motion to its seven nearest neighbors^9^. Unlike birds, the migratory cells we use for these experiments (neutrophil-like HL60 cells) cannot see, but we can effectively endow them with this sense through a microscope and a computer that tracks the motion of their neighbors. We execute seven-neighbor alignment in real time by automatically calculating the mean orientation of neighbor motion and use opto-PI3K to guide the cell in that direction (**Fig. 2A**). The cells normally lack coordination (**Fig. 2B-C, left**), but by executing this rule for each cell in the field, the cells spontaneously flock like birds (**Fig. 2B-C, middle-right, Supplemental Video 2**). Cells locally align with their neighbors, forming groups of cells that change direction together over time (**Fig. 2B, middle-right**), resulting in increasing population-level polarization (**Fig. 2C**). This classic metric of flock coordination shows that a computational rule derived from bird behavior can program collective migration *de novo* in a human immune cell line.

In simulations of bird flocking, the number of neighbors with which each individual tries to align is a key parameter that controls collective behavior^9^, but this parameter cannot be readily manipulated in living birds. With a CMI, we can vary this interaction parameter directly in living cells. We first built an agent-based simulation of the flocking rule and parameterized it to HL-60 migration speed, rotational diffusion, and response to optogenetic CMI steering. The model produced quantitatively similar total polarization over time for cells aligning with their seven closest neighbors (**Fig. 2C**). Varying the number of neighbors in the simulation predicted a decrease in polarization when cells aligned with only two neighbors and an increase when they aligned with ten (**Fig. 2D**). We then implemented these same interaction rules in living cells and observed behaviors consistent with the model predictions (**Fig. 2D, Supplemental Videos 3-4**). This experiment highlights three advantages of the CMI. First, it enables precise manipulation of interaction parameters that are difficult or impossible to perturb in natural collectives, such as the number of neighbors to which each bird responds. Second, the same interaction rule can be implanted in both simulation and in living cells, enabling direct comparison between theoretical predictions and experimental behavior. Third, changing the interaction rule requires changing only a single parameter in software, enabling rapid testing of different conditions, while holding the cellular system itself constant.

We next asked whether cellular collectives can solve guidance tasks more effectively than cells acting alone. Neutrophil-like HL-60 cells do not normally collectively migrate, so while cells close to sources of the chemoattractant peptide N-formyl-methionyl-leucyl-phenylalanine (fMLP) bias their motion toward it, distant cells are unable to interpret the direction of the gradient. To enable collective chemotactic navigation, we designed a hybrid system that combined the biologically defined gradient sensing of HL-60 with computer-defined intercellular interactions. We implemented CMI-defined cell-cell interactions with a simple rule that aligns each cell to the mean motion of the entire population (**Fig. 3A**). This rule should propagate directional information from cells that sense the fMLP gradient to the rest of the population, while the random motion of cells that do not sense the gradient should cancel out on average and contribute little to the collective direction (**Fig. 3B**). Critically, the CMI rule contains no information about the location of the source. Cells convert spatial information about the fMLP gradient into biased motion, and the CMI rule defines how that guidance information propagates through the population.

By activating the CMI interaction rule in only half of the imaging field, we could compare CMI-coordinated and control populations in the same experiment. We generated a left-to-right fMLP gradient by uncaging a photolabile fMLP analog^20,21^. Upon initiation of fMLP uncaging, both populations migrated up the resulting gradient, as demonstrated by an increase in chemotactic index (**Fig. 3C-D, Supplemental Video 5**). Activating the CMI coordination rule, however, sharply increased the chemotactic index in the CMI half of the imaging field (**Fig. 3C-D, Supplemental Video 5**). By the end of the experiment, the mean chemotactic index of the CMI-coordinated population was more than twice that of the control population (**Fig. 3D**) and remained high far from the chemotactic source (**Fig. 3E**). Thus, sharing chemotactic information through a CMI-defined interaction rule enables the population to navigate collectively, creating a capability that individual cells lack.

In the previous experiment (**Fig. 3**), guidance information was shared instantaneously across the entire population. In natural collectives, cell-cell communication is constrained in space and time by, for example, the diffusion of secreted signaling molecules. During neutrophil swarming, pioneer cells that arrive at sites of injury and infection recruit additional cells by releasing a secondary chemoattractant, the lipid LTB4. This ligand not only attracts other cells but also induces them to secrete LTB4 themselves, generating a rapidly propagating self-amplifying signal relay^5,22^(**Fig. 4A**). Our neutrophil-like HL60 cells do not normally swarm in our migration assays^23^, but we sought to reprogram them to participate in a swarm-like signal relay. Towards this end, we used the CMI to couple living cells to a simulated molecular environment. The computer calculates the production and diffusion of virtual signaling molecules in real time, while optogenetic stimulation causes the cells to migrate in response to the resulting virtual gradients. Unlike the flocking rule (**Fig. 2**) in which cells respond directly to the trajectories of their neighbors—a mode of interaction similar to vision in birds—here cells release simulated signaling molecules that form local gradients that evolve in space and time to guide the movement of surrounding cells.

We first implemented a CMI rule that enables cells to migrate up a virtual chemoattractant gradient. The CMI maintains a virtual field of chemoattractant molecules and computes diffusion in real time. At each time point, the CMI calculates the local gradient experienced by each cell and uses opto-PI3K to steer cell migration up the local gradient (**Fig. 4B**). When we introduced a constantly emitting source of virtual chemoattractant in a field of migrating HL-60 cells, this rule was sufficient to increase cell accumulation at the virtual source despite the lack of a true chemical gradient (**Fig. 4C upper, Fig. 4D, and Supplemental Video 6**). However, directional guidance was limited to cells close to the source (**Fig. 4E left**). The virtual chemotaxis rule provides a baseline for recruitment to a primary source in the absence of collective relay.

To participate in molecular relay, a cell must not only sense signals but also produce them. Inspired by the LTB4 relay that drives neutrophil swarming, we added a second CMI rule in which cells become sources of virtual chemoattractant when the local virtual chemoattractant concentration exceeds a threshold (**Fig. 4B**). Therefore, both chemoattractant production and sensing are specified computationally but coupled to the behavior of living cells to create a regenerative signal relay.

To test whether the virtual molecular relay could organize the collective recruitment of live cells, we compared cells navigating by virtual chemotaxis alone with cells using both the virtual chemotaxis and relay rules. Just as the LTB4 relay enhances the recruitment of neutrophils to primary inflammatory cues, the CMI-defined relay significantly increased cell accumulation at the primary source (**Fig. 4C lower, Fig. 4D, and Supplemental Video 7**). Pioneer cells close enough to detect the primary cue became secondary sources of virtual chemoattractant, initiating a wave of virtual signal production that propagated across the entire field of view (**Fig. 4C-D, Supplemental Video 7**), similar to the regenerative relay seen in native neutrophil swarms^5,22^. This signaling wave was accompanied by a corresponding wave of cell alignment toward the primary source **(Fig. 4E right)** recruiting cells beyond the range of the primary cue. Neither wave was observed in populations lacking the relay (**Fig. 4C lower, Fig.4D, and Fig. 4E left**).

## Discussion

Here we show that changing simple rules of cell-cell interaction is sufficient to generate new forms of collective organization and capabilities not seen in cells acting alone. We implement rules derived from systems as diverse as bird flocking and neutrophil swarming in the same cellular chassis to show that some key features of collective behavior are a function of the logic of local interactions and do not depend on the particular biological machinery that implements them. Cell-cell interactions can also extend the spatial range over which cells respond to environmental inputs, and we demonstrate two distinct strategies for long-range coordination. In one case, information is broadcast across the population; in the other, it propagates from cell to cell. Global sharing of directional information enables cells far from a chemoattractant source to benefit from information sensed by cells near the source, whereas a regenerative relay propagates this information progressively across the population through local interactions. These results show how the architecture of local interactions determines how information propagates through a collective and shapes its emergent capabilities.

The CMI provides a powerful framework for experimentally exploring these rules. Because cell-cell interaction rules are defined in software but executed by living cells, the CMI enables us to explore modes of interaction that would be difficult or impossible to implement using conventional molecular approaches, including cells that can see the motion of their neighbors. Cell-cell interaction rules can be rapidly and precisely modified in software while holding the cellular system itself constant, providing a direct way to move between theoretical models and experiments and between natural and synthetic modes of coordination in living cells. We have focused here on collective migration, but the same framework could be applied wherever individual cellular states can be measured by microscopy and dynamically controlled through light-regulated actuators. By making local interaction rules experimentally programmable, cell-machine interfaces enable an exploration of the space of possible multicellular organizations, both to uncover principles underlying natural collectives and to design collective behaviors not found in nature.

## Materials and Methods

### Cell culture

HL-60 cells came from the lab of Henry Bourne and have been checked for accuracy with STR profiling^24^. The optogenetic HL-60 line used in this work expresses a membrane anchored, light-sensitive protein, AsLov2-SsrA-mTagBFP2-CAAX, its PI3K-binding partner, ISH2-EGFP-SspBMicro, and a HaloTag-CAAX construct as described previously^19^. Undifferentiated HL-60 cells were cultured in suspensions of RPMI 1640 supplemented with L-glutamine and 25 mM HEPES (RPMI) (Corning; Corning, NY) and 10% (volume/volume) heat-inactivated fetal bovine serum (FBS) (Gibco; Waltham, MA). Cultures were sub-cultured every 2-3 days to maintain a density of 0.2–1.0 million cells/mL and incubated at 37°C/5% CO_2._ To differentiate HL-60 cells into motile, neutrophil-like cells, 1 million cells were suspended in 5 mL culture medium supplemented with 1.4% DMSO (Sigma-Aldrich) and incubated at 37°C/5% CO_2_ for 4-5 days before use in experiments.

### Preparation of cells for microscopy

Cells were imaged between a thin pad of low melting point agarose (GoldBio) and a No 1.5 glass-bottom 96-well plate (CellVis). For all experiments, the plasma membrane was visualized with JaneliaFluor JFX549^25^ conjugated to HaloTag-CAAX and nuclei were visualized in (**Figs. 2-4**) with Spy650-DNA (Spyrochrome). HL-60 cells were visually inspected to confirm differentiation and prepared for microscopy by centrifugation at 200 rcf for 10 minutes and resuspending them in RPMI containing 1 uM JaneliaFluor JFX549 and Spy650-DNA at 2x the manufacturer’s recommended concentration (Spyrochrome). After 1 hour incubating at 37°C, cells were washed in RPMI and resuspended in RPMI with 2% FBS until ready for imaging. Glass surfaces were washed by bath sonication in 5% Hellmanex III detergent before being rinsed thoroughly with water, dried, and plasma-cleaned (Harrick Plasma PDC-32G). Wells were then coated in 2% (weight/volume) bovine serum albumin (Sigma A8806) either overnight at 4° C or for 2 hours at 37° C. Agarose pads were prepared by slowly melting agarose in RPMI 1640 supplemented with L-glutamine and 25 mM HEPES at a 0.9% concentration (weight/volume) and allowed to equilibrate to 42° C before adding FBS to 2% (volume/volume). Agarose was cooled between glass slides with spacers to produce a level pad. Cells were allowed to settle to the bottom of wells for 10 minutes before agarose pads were stamped out and dropped into the well. Excess liquid was removed, compressing the cells. Except for experiments with fMLP gradients (**Fig. 3**), 10 nM fMLP was added to agarose pads to provide uniform stimulation of migration.

### Microscopy hardware

The experiments included in Figure 1 were performed on a Nikon Eclipse Ti inverted microscope equipped with a Borealis beam conditioning unit (Andor), a CSU-W1 Yokogawa spinning disk (Andor; Belfast, Northern Ireland), an iXon Ultra EMCCD camera (Andor), and a laser merge module (LMM5, Spectral Applied Research; Exton, PA) equipped with 405, 488, and 561, and 638-nm laser lines for imaging in spinning disk confocal or TIRF mode. A Nikon 20x Plan Apochromat NA 0.75 objective lens was used for (**Fig. 1A**) and a Nikon 40x Plan Apochromat NA 0.95 objective lens was used for (**Fig. 1B**). Optogenetic stimulation uses a custom DMD (Andor Technology) to spatially pattern light from an LED (470-nm) controlled by analog outputs of a digital-to-analog converter and serial commands via custom Python code. The microscope is equipped with two stacked dichroic turrets enabling simultaneous LED illumination and confocal imaging using a 488-nm long-pass dichroic filter (Chroma Technology Corp.).

All other experiments were performed on a Nikon Ti2 inverted microscope CrestOptics X-Light V3 spinning disc confocal system, a Lumencor Celesta light engine for confocal illumination, a Nikon 10x CFI Plan Apo Lambda objective, a Photometrics Kinetix sCMOS camera, and a Mightex Polygon1000 DMD, and a Lumencor Aura III LightEngine for patterned illumination with individually addressable 365nm and 488nm LEDs. The microscope is also equipped with a custom build piece of triggering hardware for sequencing multi-color optogenetic illumination^26^.

All microscope hardware was controlled using μManager^27–29^ (University of California, San Francisco) or Pycro-Manager^28^ with our custom Cell-Machine Interface platform: CyborgScope.

### Production of chemotactic gradients

For experiments with fMLP gradients (**Fig. 3**), agarose pads were supplemented with 500 nM Nv-fMLP, an fMLP derivative that is not detectable by cells until its protecting group is photochemically removed by UV illumination. The DMD was used to achieve a spatial patterning of UV light to produce a horizontal gradient. These UV light patterns were rapidly interleaved with blue light optogenetic stimulation using custom hardware^26^. The parameters for spatial patterns and the corresponding UV LED intensities are recorded in the metadata of each experiment, which will be made publicly available.

### Cell-Machine Interface platform

The CMI platform, called CyborgScope is built on two open-source software tools for computer-controlled microscopy: Pycro-Manager^28^ and μManager^27,29^. All the code used for closed-loop control of microscope hardware will be publicly available. Briefly, CyborgScope provides a flexible and generic interface for any hardware supported by μManager to operate in a closed-feedback loop according to Python code written by the user. The feedback loop consists of an imaging phase in which data is collected, and user defined image analysis software is run immediately to produce measurements of any desired cell state. Next, an optogenetic feedback control function specifies the rules by which these measurements modify the state of microscope hardware including the light-patterning hardware. Finally, the microscope hardware is updated delivering stimuli to cells which complete the loop, by changing their cell state in response to optogenetic stimuli according to biologically defined rules. Each experiment type is self-contained in a single Python file installed by the user and CyborgScope handles the image acquisition, data and metadata storage, and provides a complete record of the light stimuli delivered during the experiment. CyborgScope has a graphical user interface built in napari^30^ that allows live display of both raw images and real time image analysis results. This interface also supports user interaction via manual control of optogenetic stimuli during the experiment, if desired.

### Image segmentation, motion tracking, and optogenetic pattern generation for CMI control

Detailed methods used in each experiment are available in the acquisition class Python files within the CyborgScope codebase and the specific parameters used in each experiment can be found within the experiment metadata. The following procedure is held in common among all the CMI experiments with experiment-specific methods in the following sections. Cells were detected using signal from the HaloTag-CAAX conjugated to JFX549. Images were smoothed, thresholded according to manually selected values. Small holes were removed from the resulting binary images, and small objects were removed before producing label images. These operations were completed using the scikit-image library^31^. During the experiment, the centroids of segmented membrane signal were used to track cells using trackpy^32^. Cell-steering spots of light were positioned at the intersection point of each cell edge in the direction of desired motion with the angle of turning per frame bounded by the cells current motion vector as previously described^19^.

### CMI Cell collision algorithm

The detailed feedback control procedure for (**Fig. 1D**) can be found in crash_test_acquisition.py. Briefly, the two nearest cells in the field of view are selected to interact. Based on the current position and motion of the cells, two potential points are calculated that would cause collision at a specified angle theta, which was set to 90° for this experiment. The dot product of the vectors pointing from each cell to each point are compared and the potential collision point that is most aligned to the cells’ current motion is chosen to steer toward with optogenetic patterning as described above. This calculation is done in each frame of the time lapse, so it adjusts to variability in cell speed or how quickly cells turn in response to steering.

### CMI Cell flocking algorithm

The detailed feedback control procedure for (**Fig. 2**) can be found in cyborg_flock_acquisition.py. Each cell’s nearest n-neighbors are found using a KDTree algorithm^33^ from the SciPy library^34^ with n varying across experiments as described in the text. The mean angle of these neighbors is then used to steer cells with optogenetic patterning as described above.

### Modeling HL-60 cells for flocking simulations

Implementation of the flocking simulation is achieved using active Brownian particles with modified topological alignment modeled on a flat, two-dimensional space with periodic boundary conditions. Each particle moves at a standard speed in the direction it is oriented in. 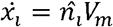. The particle experiences standard rotational diffusion, modeled with gaussian noise, but has an included term *C* in which the control signal is injected 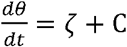, where *C* = *θ* − *θ*_*target*_. It is important to note that *C* is limited by |*C*| ≤ *C*_*max*_, where *C*_*max*_ represents the maximum turning rate that can be induced with optogenetic steering. To implement topological flocking, the desired orientation is computed using 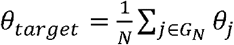 where *G*_*N*_ is group of *N* cells nearest I and in the control zone. The control signal is only active in a designated region of the simulation space and only cells within that space are considered for neighbor alignment.

The elementary active particle properties, rotational diffusion and velocity were identified by fitting the mean squared displacements of HL-60s without optogenetic steering by fitting 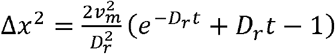 to identify *v*_*m*_ and *D*_*r*_^35^. Prior studies^19^ provided measurements of the maximum turning rates of cells during optogenetic steering which we used directly as *C*_*max*_.

### CMI Collective navigation algorithm

The detailed feedback control procedure for (**Fig. 3**) can be found in chemotactic_hive_mind_acquisition.py. In this experiment, a pre-determined set of instructions is generated for the UV photo stimulation that generates the chemotactic gradient of fMLP. These instructions are interleaved with blue light optogenetic stimulation using custom hardware^26^ to rapidly switch between UV light patterns that shape the gradient, and blue light patterns that optogenetically steer cell motion as described above. The mean motion angle of all cells that are migrating within the CMI region of the field of view is calculated and this angle is used to steer each cell as described above. Note that cells in the negative control area (No CMI) do not contribute to the collective sensing of the fMLP gradient.

### Modeling Virtual Chemical Cues, Diffusive Relays and the CMI Swarming algorithm

The spatiotemporal dynamics of the virtual chemical signal *c*(*x, t*) are governed by a two-dimensional reaction-diffusion equation:

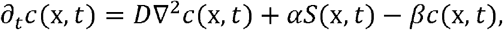

where *D* is the cue diffusion coefficient, *β* is the degradation rate, *α* is the chemical secretion rate density, and *S*(*x, t*) ∈ [0, 1] represents the spatial source term. Absorbing Dirichlet boundary conditions (*c* = 0) are enforced at the perimeter of the field of view. To achieve real-time feedback during image acquisition, the diffusion term is integrated using an implicit backward Euler scheme, pre-factorized via sparse LU decomposition, while reaction and decay are handled explicitly. The PDE is solved on a coarse spatial grid (downsampled by a factor of 5 in comparison with the microscope resolution) and bilinearly interpolated back to camera resolution at each time step.

Individual cells interact with this virtual field through their two-dimensional footprints. Rather than evaluating point values or spatial derivatives, each cell senses the local cue by integrating and averaging the concentration over its mask. Directional sensing for closed-loop optical steering is achieved by calculating the intensity-weighted center of mass of the chemical field across the cell’s mask; the vector pointing from the cell centroid to this weighted center defines the target heading, prompting the projector to position a localized light stimulus at the cell’s leading edge.

In the chemotaxis experiments (**Fig. 4**), both with and without relay, the primary cue is supplied by a constant external source at the center of the field of view, which adds a fixed amount of the virtual chemical to a 25 × 25 pixel square (17 × 17 *µ*m^2^) at every frame.

For diffusive relay experiments, the source term *S*(*x, t*) is generated by the cells themselves following the threshold-activated relay model of Dieterle et al^22^. Here, cells act as excitable signal repeaters: when the mean concentration across a cell’s mask exceeds an activation threshold *C*_th_, the cell switches irreversibly to an active relay state and secretes the virtual chemical across its footprint (*S* = 1) from the next frame onward. To prevent stationary cells or segmentation debris from becoming relays, activation additionally requires that the cell moved by more than 2.5 camera pixels (1.7 *µ*m) since the previous frame. Independently of their relay state, cells are optically steered up the local gradient whenever their mean signal exceeds a lower detection threshold *C*_det_.

The relay experiments used *D* = 125 *μ*m^2^/s, *α* = 0.03 nM/s, *β* = 0.005 s^−1^, *C*_det_ = 0.01 nM and *C*_th_ = 0.05 nM.

Without relay, degradation confines the cue to the vicinity of the primary source: on the timescale 1/β ≈ 3 min, the field reaches a steady state that extends over the decay length 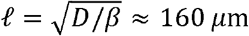 and falls off exponentially beyond it. With relay, the activation front, i.e. the largest distance from the source at which at least half of the cells have become relays, reached ~ 1 mm, about six decay lengths (**Fig. 4**).

### Image analysis

For quantification of cell motion in (**Figs. 2-4**), cell position was tracked by nuclear position instead of using the centroid of membrane signal because nuclei are less prone to collision and enable more accurate tracking. All analysis code and output data will be made available upon final publication. Briefly, after median filtering, nuclei were detected using a Laplacian of Gaussian filter^31,36^ using manually-selected parameters and assigned to tracks with trackpy^32^. Tracks were visually inspected for accuracy by plotting them over the raw images.

### Population Polarization

Population polarization can be computed for each frame of simulation or experimental data. This scalar metric is the magnitude of the average orientation vectors of all particles at a given timepoint.

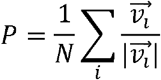

The population polarization provides an evaluation of the cluster’s motion as a whole and is useful for identifying state transitions ^12^ as few cells are required to generate a meaningful measurement. Averages of this metric are taken across timepoints and simulations without weighting.

### Correlation Length

Correlation length quantifies the scale of alignment between individual cells in simulation. It is computed on the motion of individual cells relative to the entire swarm.

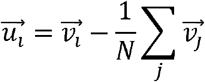

A correlation function C(r) is computed as follows, where r describes the distance between each pair of particles.

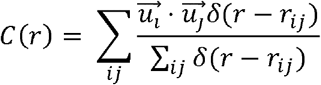

C(r) has been found to be well described by a power law ^37^.

To compute the specific correlation distance. We computer C(R) and perform logarithmic binning. As this is the correlation of velocity fluctuations, the correlation function is expected to be negative at sufficiently long ranges. We use the fitted power law to identify the distance at which the correlations in velocity fluctuations have fallen to zero.

Accumulating sufficient statistics to fit C(r) well typically exceeds the quantity of data available in a single simulation or experimental timepoint. To merge experimental data all the combined C(r) points collected from a steady state are joined into one then are binned and fit together. Handling extremely large data volumes benefits mitigating the n^2 scaling of the C(r) data set. To accomplish this, the data is binned first and the binned data from the steady state and replicate simulations are merged before finally being fit.

The correlation length metric is unlike population polarization which only describes collective as a whole and does not take into account the relationships between nearby cells. The final measurement produces a real length beyond which cells are on average do not respond to each other.

Chemotactic index was calculated as the cosine of the angle between cell motion vector and either a horizontal vector in (**Fig. 3**) or the vector from the cell nucleus to the simulated chemotactic source (**Fig. 4**). The parameters for binning motion in space and time can be found in the code used to generate figures.

## Resource availability

### Data Availability

Upon final publication, uncropped versions of microscopy images used in figures, as well as all tracking data, and metadata will be deposited at Zenodo and made publicly available. The cell line used in this work and raw microscopy data is available upon request from O.D.W.

### Code Availability

Upon final publication all code used for data analysis, figure generation, and microscopy hardware control will be made publicly available at Zenodo. Any additional information required to reanalyze the data reported in this paper is available from the lead author upon request.

## Acknowledgements

O.D.W. is funded by the National Institutes of Health grant GM118167, the National Science Foundation Center for Cellular Construction grant DBI-1548297 and the Sandler Program for Breakthrough Biomedical Research, which is partially funded by the Sandler Foundation. N.R.M. is supported by a Cancer Research Institute Irvington Postdoctoral Fellowship. N.B. was supported by a National Institutes of Health, NIH G-RISE, T32GM141862 Fellowship (PI - A.G.) A.G also acknowledges support from the National Science Foundation, NSF-CREST: Center for Cellular and Biomolecular Machines at UC Merced, NSF-HRD-2112675 and the National Science Foundation, Center for Engineering Mechanobiology, grant CMMI-1548571. A.A. thanks the generous support of the Helen and Martin Kimmel Award for Innovative Investigation and the Clore Center for Biological Physics. Photocaged fMLP was a generous gift from Elizabeth Sisko and the lab of Jason Sello. JFX549 HaloTag dye was kindly provided by Dr. Luke Lavis. N.M. thanks Jason P. Town and Nico Stuurman for their help in implementing computer control of microscope hardware and their inspiration to pursue ever more complex forms thereof. Thanks to Alex Janzen at Mightex for his assistance in maximizing the use of our DMD hardware. Thanks to Evelyn D. Strickland for sharing her expertise in neutrophil swarming and for her lively discussions. Finally, thanks to Henry R. Scott for sharing his enthusiasm for, and belief in this project from its inception.

## Figures and Video Captions

**Supplemental Video 1: Programmed Alignment of Cell Motion Leads to Collisions**. HL-60 neutrophil-like cells can be steered by automated segmentation and optogenetic stimulation of opto-PI3K^19^. The optogenetic illumination pattern is visualized in magenta. First, a user-defined command steers the cell to the top right in the image reference frame. Second, the CMI enables optogenetic stimuli that depend on the behavior of the cells themselves rather than a human user. Here, stimulation of one cell adjusts to the motion of the other such that they align at 90° to one another, eventually colliding. Neither the location nor the path to collision are prespecified, and these depend entirely on the starting motility of the cells and their responses to each other mediated by the CMI.

**Supplemental Video 2: Cell flocking with a 7-neighbor alignment rule**. HL-60 neutrophil-like cells exhibit random uncoordinated motion in the absence of external chemotactic gradients or optogenetic stimulation. A CMI-defined 7-neighbor alignment rule is enabled after approximately 16 minutes, leading to cell alignment and locally correlated changes in motion just like bird flocks. Cell track segments are color-coded by angle. Scale bar is 500 μm. Images were taken every 10 seconds.

**Supplemental Video 3: Cell flocking with a 2-neighbor alignment rule**. HL-60 neutrophil-like cells exhibit random uncoordinated motion in the absence of external chemotactic gradients or optogenetic stimulation. A CMI-defined 2-neighbor alignment rule is enabled after approximately 15 minutes, leading to cell alignment and locally correlated changes on shorter length scales than the 7-neighbor rule. Cell track segments are color-coded by angle. Scale bar is 500 μm. Images were taken every 10 seconds.

**Supplemental Video 4: Cell flocking with a 10-neighbor alignment rule**. HL-60 neutrophil-like cells exhibit random uncoordinated motion in the absence of external chemotactic gradients or optogenetic stimulation. A CMI-defined 10-neighbor alignment rule is enabled after approximately 15 minutes, leading to cell alignment and locally correlated changes on longer length scales than the 7-neighbor rule. Cell track segments are color-coded by angle. Scale bar is 500μm. Images were taken every 10 seconds.

**Supplemental Video 5: Global coordination enhances collective chemotactic guidance**. A Chemotactic gradient is introduced after approximately 16 minutes, and the global coordination CMI rule was enabled in the top half of the field after an additional 8 minutes, improving the ability of the coordinated cells to migrate towards the source compared to control cells. Scale bar is 250μm.

**Supplemental Video 6: Passive diffusion of a virtual chemoattractant produces short-range recruitment**. HL-60 neutrophil-like cells (green) migrate in the absence of a physical chemotactic gradient and are guided by a simulated diffusible chemoattractant (magma colormap) using a CMI “chemotaxis-only” rule. A constant source of virtual chemoattractant is placed near the center of the field generates a gradient that evolves by passive diffusion. Cells locally climb this virtual gradient and accumulate near the source, but guidance is limited to cells within a short range of the source. Scale bar is 500 μm. Images were taken every 10 seconds.

**Supplemental Video 7: Synthetic signal relays propagate guidance information to enable long-range recruitment**. HL-60 neutrophil-like cells (green) migrate in the absence of a physical chemotactic gradient are guided by a simulated diffusive chemoattractant (magma colormap) using a CMI “chemotaxis+signal relay” rule. As in Video 6, a constant source of virtual chemoattractant near the center of the field produces a gradient by passive diffusion. Here, however, cells that encounter a sufficiently high concentration of the virtual chemoattractant become secondary sources themselves, producing a regenerative signal relay. The resulting wave of virtual chemoattractant propagates across the field, extending guidance beyond the range of the primary source, and recruiting cells from greater distances. Scale bar is 500 μm. Images were taken every 10 seconds.

## Bibliography

1. Vicsek, T. & Zafeiris, A. Collective motion. Phys. Rep. 517, 71–140 (2012).

2. David Morgan, E. Trail pheromones of ants. Physiol. Entomol. 34, 1–17 (2009).

3. Gordon, D. M. The Ecology of Collective Behavior in Ants. Annu. Rev. Entomol. 64, 35–50 (2019).

4. Chtanova, T. et al. Dynamics of neutrophil migration in lymph nodes during infection. Immunity 29, 487–496 (2008).

5. Strickland, E. et al. Self-extinguishing relay waves enable homeostatic control of human neutrophil swarming. Dev. Cell 59, 2659–2671.e4 (2024).

6. Lämmermann, T. et al. Neutrophil swarms require LTB4 and integrins at sites of cell death in vivo. Nature 498, 371–375 (2013).

7. Hino, N. et al. ERK-Mediated Mechanochemical Waves Direct Collective Cell Polarization. Dev. Cell 53, 646–660.e8 (2020).

8. Scarpa, E. & Mayor, R. Collective cell migration in development. J. Cell Biol. 212, 143–155 (2016).

9. Ballerini, M. et al. Interaction ruling animal collective behavior depends on topological rather than metric distance: Evidence from a field study. Proc. Natl. Acad. Sci. 105, 1232–1237 (2008).

10. Buhl, C. et al. From Disorder to Order in Marching Locusts. Science 312, 1402–1406 (2006).

11. Partridge, B. L. & Pitcher, T. J. The sensory basis of fish schools: Relative roles of lateral line and vision. J. Comp. Physiol. 135, 315–325 (1980).

12. Vicsek, T., Czirók, A., Ben-Jacob, E., Cohen, I. & Shochet, O. Novel Type of Phase Transition in a System of Self-Driven Particles. Phys. Rev. Lett. 75, 1226–1229 (1995).

13. Collective behavior from surprise minimization | PNAS. https://www.pnas.org/doi/10.1073/pnas.2320239121 (2025).

14. Stricker, J. et al. A fast, robust and tunable synthetic gene oscillator. Nature 456, 516–519 (2008).

15. Elowitz, M. B. & Leibler, S. A synthetic oscillatory network of transcriptional regulators. Nature 403, 335–338 (2000).

16. Gardner, T. S., Cantor, C. R. & Collins, J. J. Construction of a genetic toggle switch in Escherichia coli. Nature 403, 339–342 (2000).

17. Basu, S., Gerchman, Y., Collins, C. H., Arnold, F. H. & Weiss, R. A synthetic multicellular system for programmed pattern formation. Nature 434, 1130–1134 (2005).

18. Sprinzak, D. & Elowitz, M. B. Reconstruction of genetic circuits. Nature 438, 443–448 (2005).

19. Town, J. P. & Weiner, O. D. Local negative feedback of Rac activity at the leading edge underlies a pilot pseudopod-like program for amoeboid cell guidance. PLOS Biol. 21, e3002307 (2023).

20. Pirrung, M. C., Drabik, S. J., Ahamed, J. & Ali, H. Caged chemotactic peptides. Bioconjug. Chem. 11, 679–681 (2000).

21. Collins, S. R. et al. Using light to shape chemical gradients for parallel and automated analysis of chemotaxis. Mol. Syst. Biol. 11, 804 (2015).

22. Dieterle, P. B., Min, J., Irimia, D. & Amir, A. Dynamics of diffusive cell signaling relays. eLife 9, e61771 (2020).

23. Babatunde, K. A. et al. Chemotaxis and swarming in differentiated HL-60 neutrophil-like cells. Sci. Rep. 11, 778 (2021).

24. Saha, S., Town, J. P., Weiner, O. D. & Huttenlocher, A. Mechanosensitive mTORC2 independently coordinates leading and trailing edge polarity programs during neutrophil migration. Mol. Biol. Cell 34, ar35 (2023).

25. Grimm, J. B. et al. A general method to optimize and functionalize red-shifted rhodamine dyes. Nat. Methods 17, 815–821 (2020).

26. Martin, N. nicmarti4801/OptoTrigger: v1.0.0. https://doi.org/10.5281/zenodo.21283606 (2026) doi:10.5281/zenodo.21283606.

27. Edelstein, A., Amodaj, N., Hoover, K., Vale, R. & Stuurman, N. Computer Control of Microscopes Using µManager. Curr. Protoc. Mol. Biol. 92, 14.20.1-14.20.17 (2010).

28. Pinkard, H. et al. Pycro-Manager: open-source software for customized and reproducible microscope control. Nat. Methods 18, 226–228 (2021).

29. Edelstein, A. D. et al. Advanced methods of microscope control using μManager software. J. Biol. Methods 1, 1 (2014).

30. Sofroniew, N. et al. napari: a multi-dimensional image viewer for Python. https://doi.org/10.5281/zenodo.7276432 (2022) doi:10.5281/zenodo.7276432.

31. Walt, S. van der et al. scikit-image: image processing in Python. PeerJ 2, e453 (2014).

32. Allan, D. B., Caswell, T., Keim, N. C., van der Wel, C. M. & Verweij, R. W. soft-matter/trackpy: v0.7. https://doi.org/10.5281/zenodo.16089574 (2025) doi:10.5281/zenodo.16089574.

33. Maneewongvatana, S. & Mount, D. M. Analysis of approximate nearest neighbor searching with clustered point sets. Preprint at 10.48550/arXiv.cs/9901013 (1999).

34. Virtanen, P. et al. SciPy 1.0: fundamental algorithms for scientific computing in Python. Nat. Methods 17, 261–272 (2020).

35. Fürth, R. Die Brownsche Bewegung bei Berücksichtigung einer Persistenz der Bewegungsrichtung. Mit Anwendungen auf die Bewegung lebender Infusorien. Z. Phys. 2, 244–256 (1920).

36. Otero, I. R. & Delbracio, M. Anatomy of the SIFT Method. Image Process. Line 4, 370–396 (2014).

37. Cavagna, A. et al. Scale-free correlations in starling flocks. Proc. Natl. Acad. Sci. 107, 11865–11870 (2010).

